# Microbial Reductive Dechlorination of Hexachlorobutadiene by *Dehalococcoides*

**DOI:** 10.64898/2026.10.15.759271

**Authors:** Xin Wang, Hongyan Wang, Xinyue Qian, Haiwei Wei, Yin Zhong, Zongming Xiu, Haiyan Zhang, Lian Yu, Shanquan Wang, Haixia Yan, Wenjing Sun, Yi Yang

## Abstract

Hexachlorobutadiene (HCBD), a persistent organic pollutant (POP), poses a significant threat to ecological environments and human health. While anaerobic microbial transformation is a promising remediation strategy, the specific microorganisms and metabolic mechanisms involved have remained poorly understood. This study demonstrates the successful enrichment of an anaerobic microbial consortium from river sediment, with lactate as an electron donor, capable of transforming HCBD. Within 240 days, 62.5 μmol of HCBD was sequentially dechlorinated through pentachlorobutadiene (PCBD), tetrachlorobutadiene (TeCBD), trichlorobutadiene (TCBD), and dichlorobutadiene (DCBD) to chlorobutadiene (CBD). Co-amendment with HCBD and trichloroethene (TCE) resulted in complete removal of both compounds. 16S rRNA gene sequencing and quantitative PCR showed enrichment of *Dehalococcoides*, linking its population growth to HCBD and TCE dechlorination. This study demonstrates that *Dehalococcoides* growth was associated with HCBD transformation in the enrichment cultures, broadens our understanding of the metabolic capabilities of *Dehalococcoides*, and identifies a candidate gene for this important environmental process, offering valuable insights for the bioremediation of sites contaminated with HCBD.

## Introduction

Hexachlorobutadiene (HCBD) is a synthetic chemical historically produced for commercial applications, notably as a solvent, and is also an unintentional byproduct in the manufacturing of chlorinated hydrocarbons such as trichloroethene (1, 2). With no known natural sources, its presence in the environment is due solely to human activity (3, 4). Once released from industrial or waste sites into aquatic environments, HCBD’s pronounced hydrophobicity (e.g., low aqueous solubility) causes it to partition from water and accumulate in sediments and soils (5, 6). This creates persistent, long-term contaminant reservoirs that are challenging to remediate and pose a significant environmental concern (7). Despite a low vapor pressure, HCBD can volatilize from contaminated surfaces over time. Once in the atmosphere, it is highly persistent, with an estimated half-life ranging from 60 days to over 3 years due to slow oxidation by hydroxyl radicals (8). This longevity enables long-range atmospheric transport, resulting in widespread pollution far from its industrial origins. Consequently, HCBD is now a ubiquitous contaminant, detected in diverse environmental media—including air, water, soil, and sediment—across numerous countries and has even been found in remote ecosystems like the Arctic (9–11). Its detection in biota, from aquatic life to Arctic mammals and birds, confirms its ability to enter and bioaccumulate in global food webs. Recognized for its high toxicity to aquatic life, HCBD is also a probable human carcinogen that causes severe, targeted kidney damage and is genotoxic (12). In recognition of its combined toxicity, persistence, bioaccumulation, and capacity for long-range transport, the Stockholm Convention officially listed HCBD as a persistent organic pollutant (POP) in 2015 (13), mandating its global restriction and elimination. Therefore, HCBD is a contaminant of global concern, demanding a comprehensive understanding of its environmental fate to mitigate its risks.

The development of effective bioremediation strategies for HCBD is contingent upon a thorough understanding of its microbial transformations. Aerobic HCBD removal accompanied by chloride release has been reported for *Serratia marcescens* strain HL1 (14), and anaerobic reductive dechlorination represents another promising pathway for HCBD biotransformation (15). However, knowledge of the microorganisms responsible for anaerobic HCBD dechlorination and their underlying dechlorination mechanisms remains limited. Initial studies demonstrated the feasibility of this pathway, with sediment cultures and methanogenic enrichments converting HCBD into various partially dechlorinated products, including tetrachloro-, trichloro-, and dichloro-butadiene isomers (16, 17). To date, only a single study has begun to elucidate the role of a specific microorganism in this process. *Dehalococcoides*, a known organohalide-respiring bacterium, was shown to co-metabolically dechlorinate HCBD to (E)-1,2,3-trichlorobutadiene [(E)-1,2,3-tri-CBD] via sequential intermediates, including (E)-1,1,2,3,4-pentachlorobutadiene [(E)-1,1,2,3,4-penta-CBD] and (Z,E)-1,2,3,4-tetrachlorobutadiene [(Z,E)-1,2,3,4-tetra-CBD] when simultaneously growing on tetrachloroethene (18). However, this process did not support microbial growth and the dechlorination remained incomplete. While theoretical calculations suggest complete dechlorination is possible (18), and some non-specific reactions using cyanocobalamin can yield chlorine-free products, the specific microorganisms and pathways remain unknown (19, 20). This highlights some critical gaps in the current knowledge: (1) no microorganism has been enriched and/or isolated that can utilize HCBD as a sole electron acceptor for growth, (2) the metabolic pathway from HCBD to non-chlorinated compound has not been documented, and (3) the key enzymes responsible for a complete dechlorination sequence are unidentified. Addressing these questions is crucial to understanding the environmental fate of HCBD and developing targeted microbial solutions for its removal. This study presents characterization of a novel *Dehalococcoides* sp. strain HCBD that utilizes hexachlorobutadiene (HCBD) as a growth-supporting electron acceptor under anoxic conditions. Originating from river sediment, this enrichment provides a crucial advancement, as *Dehalococcoides* is a genus renowned for its vital role in the detoxification of organohalides (21). We offer multiple lines of evidence for this discovery, including monitoring dechlorination activity and quantifying population growth via qPCR, which showed an association between cell proliferation and HCBD transformation. Furthermore, we proposed a reductive dechlorination pathway based on the detected transformation products of HCBD and identified a putative KB1-5 like reductive dehalogenase (99.60% similarity) through metagenomics (22), with its expression supported by proteomic detection. The presence of this KB1-5-like enzyme likely explains the consortium’s observed ability to transform both HCBD and trichloroethene (TCE). Collectively, these results substantially broaden our understanding of the metabolic diversity and substrate range of *Dehalococcoides*, representing a significant advance for microbial ecology and the development of effective bioremediation technologies for HCBD.

## Results

### Reductive Dechlorination of HCBD under Anoxic Conditions

Following an extended eight-month incubation, microcosms containing sediment from the Xi River demonstrated the sequential reductive dechlorination of hexachlorobutadiene (HCBD) to 2-chlorobuta-1,3-diene (CBD) (Figure 1A). The biotransformation of the initial 62.5 μmol of HCBD proceeded in a stepwise manner, with the pathway elucidated using enrichment cultures and product identification by GC-MS (Table S2 and Figures S5-S10). During the first 30 days of incubation, HCBD concentration gradually decreased as the primary intermediates, 1,1,2,4,4-pentachlorobuta-1,3-diene (PCBD) and (3Z)-1,1,3,4-tetrachlorobuta-1,3-diene (TeCBD), appeared. Subsequently, after approximately 60 days, the lesser-chlorinated intermediates (E)-1,2,4-trichlorobuta-1,3-diene (TCBD) and 1,2-dichlorobuta-1,3-diene (DCBD) were detected. The final product, CBD, began to manifest and was detected until the end of the 240-day incubation period (Figure 1B). At the end of the experiment on day 240, HCBD and PCBD were no longer detectable, while TeCBD, TCBD, DCBD, and CBD persisted as the daughter products of the observed transformation.

**Figure 1.**
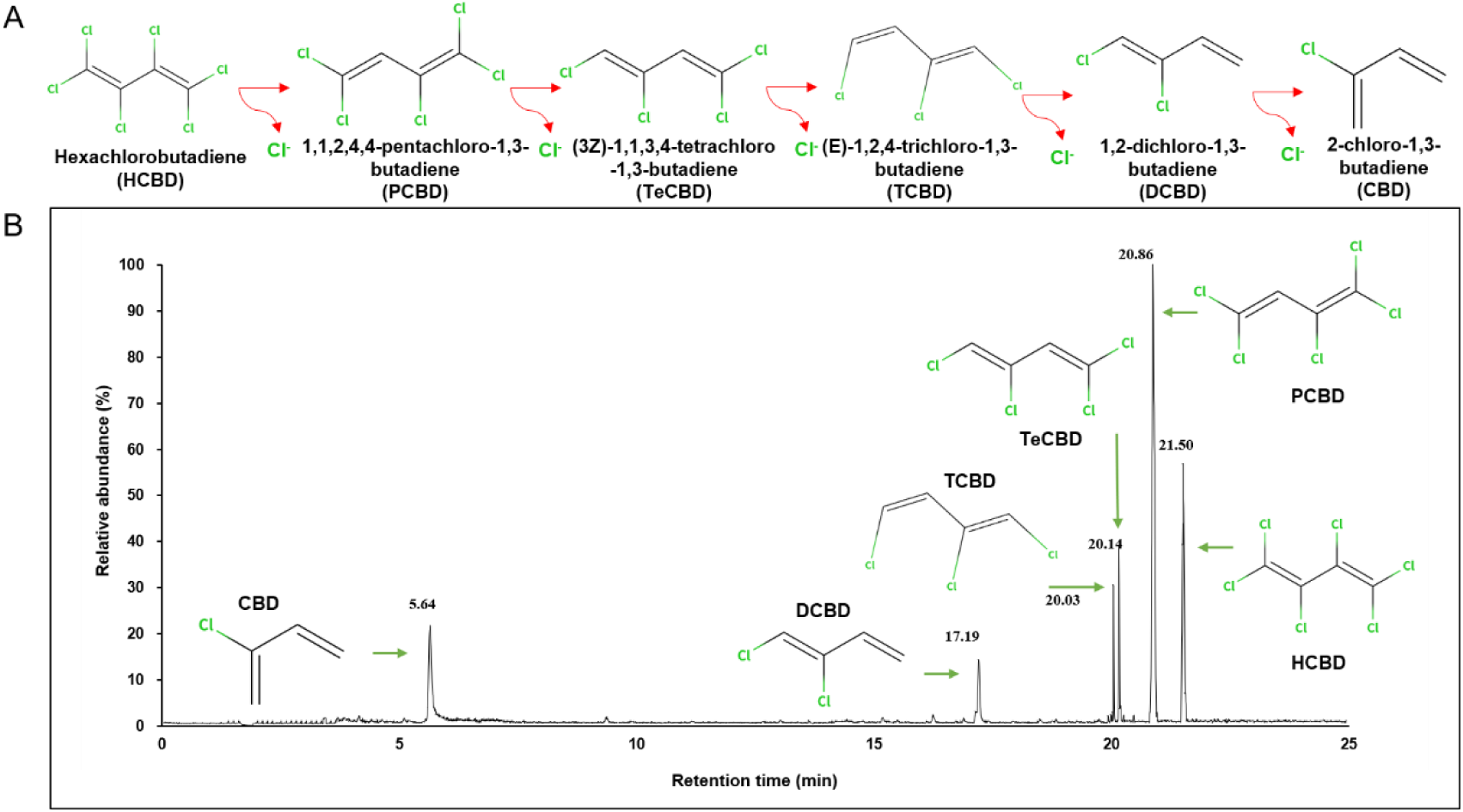
Panel A shows the sequential reductive dechlorination proposed pathway of HCBD under anoxic conditions. Each transformation involves Cl^−^ removal, progressing from HCBD → 1,1,2,4,4-PCBD → (3Z)-1,1,3,4-TeCBD → (E)-1,2,4-TCBD → DCBD → CBD. Panel B displays the GC-MS chromatogram showing relative abundance with retention times on day100: CBD (5.64 min), 1,2-DCBD (17.19 min), (E)-1,2,4-TCBD (20.03 min), (3Z)-1,1,3,4-TeCBD (20.14 min), 1,1,2,4,4-PCBD (20.86 min), and HCBD (21.50 min).

### Microbial Dechlorination of TCE and HCBD Simultaneously

The capacity of the HCBD-enriched culture to dechlorinate TCE was examined. The less-acclimated, first transfer culture fed with HCBD and TCE (T1-HCBD+TCE) achieved complete transformation of TCE to 80.5 μmol of ethene by day 81 (Figure 2A). In the third-transfer culture fed only with TCE (T3-TCE), 81 μmol of TCE was depleted within 80 days, stalling at *cis*-DCE (46 μmol) and producing methane (15 μmol) (Figure 2C). In contrast, cultures amended with both HCBD and TCE showed a shift towards complete dechlorination of TCE to ethene (ETH). The third transfer culture of HCBD and TCE (T3-HCBD+TCE) converted the TCE to 48 μmol of ethene after 95 days, with no methane production (Figure 2B). The presence of TCE also influenced the extent of HCBD dechlorination. The most extensive dechlorination of HCBD occurred in the T1-HCBD+TCE culture, which produced DCBD, while the T3 cultures (T3-HCBD) showed less extensive degradation, yielding primarily TCBD and TeCBD.

**Figure 2.**
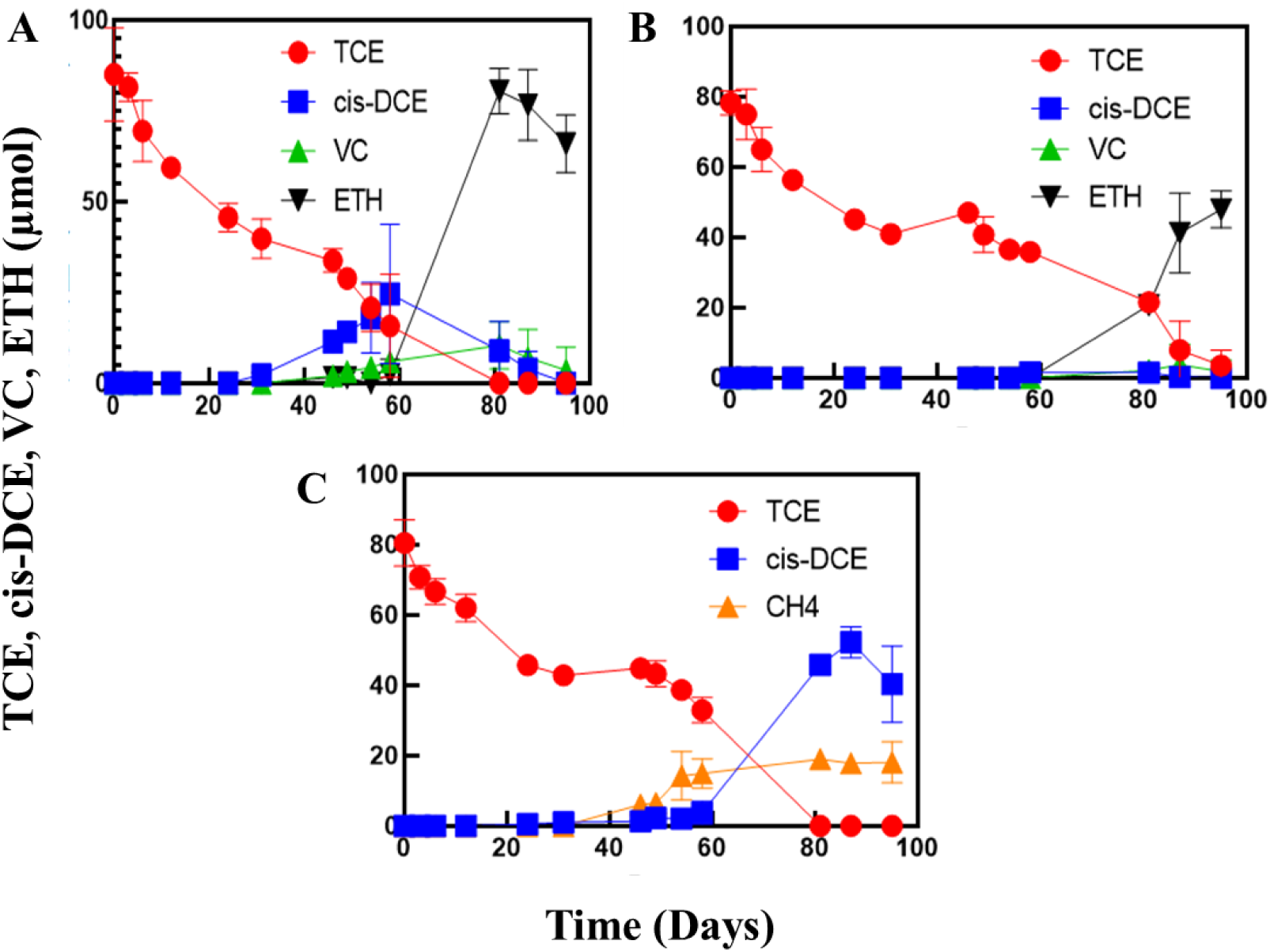
Reductive dechlorination of TCE under different enrichment conditions (n=3). (A) TCE degradation in T1-HCBD+TCE system showing complete transformation of TCE to ethene with intermediate formation of cis-DCE and VC. (B) TCE degradation in T3-HCBD+TCE system demonstrating conversion to ethene with minimal intermediate accumulation. (C) TCE degradation in T3-TCE system showing incomplete dechlorination with cis-DCE as terminal products.

### Enrichment of *Dehalococcoides* during dechlorination

Amplicon sequencing was used to track microbial community shifts from the initial river sediment through microcosm enrichment and subsequent transfers. The community composition changed from the sediment to the enrichment cultures (Figure 3A). *Proteobacteria*, initially the most abundant phylum (34.3%), decreased sharply to 3.4% in the microcosm before rebounding in later transfers. The relative abundance of *Chloroflexi*, the phylum containing *Dehalococcoides*, was stable initially (15.34%) before decreasing in the second transfer. Other phyla like *Cloacimonadota*, *Spirochaetota*, and *Thermotogota* remained present at low, stable abundances throughout the enrichment process. A detailed genus-level analysis highlighted the selection process (Figure 3B). The initial microcosm cultivation prompted a bloom of certain genera, notably *Christensenellaceae* R-7, which surged from just 0.09% in the sediment to become the most abundant genus at 11.3%. However, this dominance was transient, as its abundance plummeted to less than 1% after subsequent transfers. A similar pattern was observed for *Cryptanaerobacter*. In contrast, *Dehalococcoides*, the key organohalide-respiring bacterium, underwent a significant and targeted enrichment. Its relative abundance increased more than 50-fold, from 0.2% in the original sediment to 10.7% in the initial microcosm where active HCBD degradation was occurring. While its abundance fluctuated in subsequent transfers (4.6% and 1.2%), this initial, sharp increase strongly links *Dehalococcoides* to the observed dechlorination activity. Consistent with these compositional shifts, α-diversity metrics revealed that the HCBD+TCE microcosm harbored the most diverse and even microbial community (ACE = 2045.39; Chao1 = 2030.79; Shannon = 7.93; Simpson = 0.982), exceeding that of the HCBD-only microcosm (ACE = 1625.93; Chao1 = 1558.62; Shannon = 6.38; Simpson = 0.957). Community richness and diversity progressively changed upon serial transfers, with T1 (ACE = 1131.00; Chao1 = 1094.34; Shannon = 5.36; Simpson = 0.901) and, more markedly, T2 (ACE = 341.40; Chao1 = 325.93; Shannon = 5.17; Simpson = 0.942) exhibiting stepwise change (Table S5). Collectively, these patterns indicate that successive subculturing under sustained selective pressure constrained overall community breadth while preserving the core functional guilds responsible for reductive dechlorination, particularly *Dehalococcoides*.

**Figure 3.**
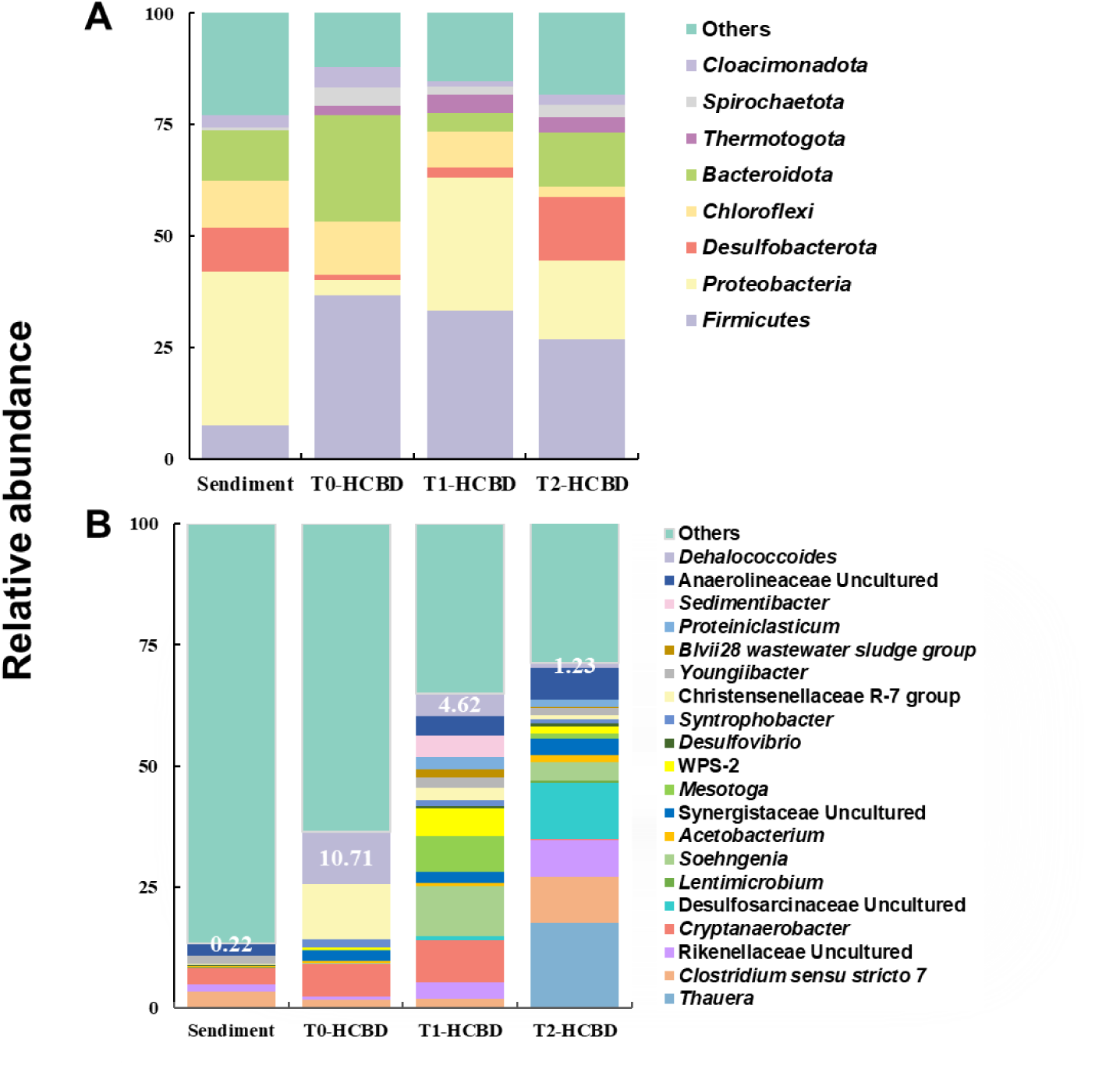
Microbial community structures at the phylum level (A) and the genus level (B) in the river sediment, HCBD-dechlorinating sediment microcosms and enrichment transfer cultures.

### Growth of *Dehalococcoides* during HCBD reductive dechlorination

To confirm that *Dehalococcoides* growth was coupled with HCBD, we quantified its cell density using qPCR across various enrichment stages and substrate conditions (Figure S4). In cultures fed only with HCBD, *Dehalococcoides* consistently showed growth linked to dechlorination. The population increased 37.5-fold in the initial microcosm (T0-HCBD), and this growth was sustained, albeit at lower levels (7.9-fold and 7.0-fold), through the first two transfers. The presence of trichloroethene (TCE) alongside HCBD, however, resulted in robust growth (144.9-fold; T0-HCBD+TCE).

### Phylogenetic Placement of the Enriched *Dehalococcoides* sp. strain HCBD

To identify the specific *Dehalococcoides* population responsible for HCBD dechlorination, we amplified, sequenced, and phylogenetically analyzed a 1301 bp fragment of its 16S rRNA gene recovered from the metagenome-assembled genome. The resulting sequence, designated *Dehalococcoides* sp. strain HCBD, was compared with known organohalide-respiring bacteria (Figure 4, Figure S3). Phylogenetic analysis revealed that strain HCBD is closely related to a specific clade of *Dehalococcoides mccartyi*, showing the sequence similarity (99.85%) to strains FL2, CBDB1, 11a5, and JNA. Its similarity to other well-known *Dehalococcoides* strains, including BAV1, CG3, VS, and 195, was also high, ranging from 90% to 99.93%. In contrast, its similarity to strains from the more distantly related *Dehalogenimonas* genus was lower (88.12% to 90.28%).

**Figure 4.**
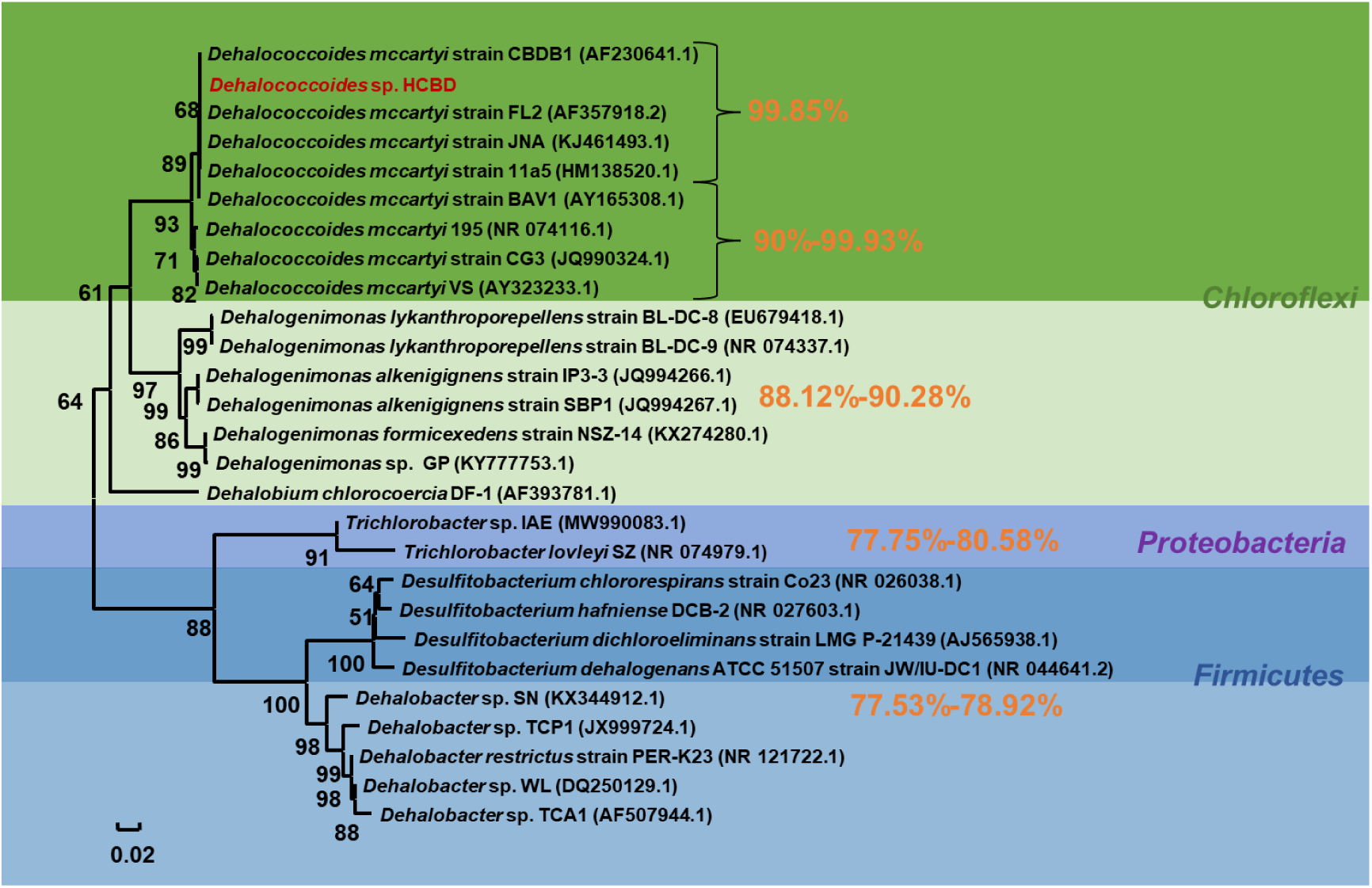
Phylogenetic tree of *Dehalococcoides* sp. HCBD (red highlight) from HCBD-dechlorinating enrichment cultures based on the nearly full-length 16S rRNA genes. Numbers beside branches indicate percentage of branch support based on 1000 bootstrap re-sampling. The scale bar indicates an evolutionary distance of 0.02 nucleotide substitutions per site.

### Metagenomic Profile of the HCBD-Dechlorinating Enrichment Culture

Genomic DNA from the second transfer of an enrichment culture, utilizing lactate as an electron donor and carbon source and HCBD as an electron acceptor, was subjected to metagenomic sequencing. From 69,713,006 original sequence reads, 49,969,086 high-quality sequences (72%) were retained after quality control. Microbial community composition, excluding taxa with relative abundances below 0.5%, was analyzed at phylum, class, and genus levels (Figure S1). At the phylum level, eight dominant phyla were identified: *Euryarchaeota* (1,746,421 sequences, 65.2%), *Proteobacteria*, *Firmicutes*, *Bacteroidetes*, *Chloroflexi*, *Spirochaetes*, *Actinobacteria*, and *Synergistetes*. *Proteobacteria*, *Firmicutes*, and *Chloroflexi*, known to include organohalide-respiring bacteria, collectively accounted for 577,804 sequences (21.6%). At the class level, eight classes predominated: *Methanomicrobia* (1,743,420 sequences, 65%), *Clostridia*, *Deltaproteobacteria*, *Betaproteobacteria*, *Bacteroidia*, *Dehalococcoidia* (61,518 sequences, 2.3%), *Bacilli*, and *Spirochaetia*. At the genus level, 11 genera were prominent: *Dehalococcoides* (60,938 sequences, 2.3%), *Methanosarcina*, *Methanocorpusculum*, *Thauera*, *Desulfovibrio*, *Pelotomaculum*, *Lentimicrobium*, *Methanofollis*, *Sphaerochaeta*, *Petrimonas*, and *Methanoculleus*. The significant presence of *Dehalococcoides* underscores its role in HCBD reductive dechlorination.

### Draft Genome Analysis of strain HCBD

Metagenomic assembly and binning of the enrichment culture yielded 26 high quality genome bins, including 21 bacterial and 5 archaeal bins (Figure S1). Among these, a near-complete draft genome of a *Dehalococcoides* strain designated *Dehalococcoides* sp. strain HCBD was recovered, with an estimated completeness of 99.01% and 0% contamination. The genome spans 1,227,134 bp across 17 contigs, with an N50 of 77,826 bp, L50 of 3, and includes 48 RNA features. Species identification using the TYGS indicated a DDH value of 73.6% (confidence interval: 69.7– 77.3%) relative to *Dehalococcoides mccartyi* strain 195^T^, suggesting that strain HCBD represents a new member within the *Dehalococcoides mccartyi* species. Whole-genome phylogenetic analysis (Figure S2) confirmed that strain HCBD clusters most closely with *Dehalococcoides mccartyi* 195^T^. The overall genomic features of strain HCBD are comparable to those of reference strains *Dehalococcoides mccartyi* CBDB1 and 195 ^T^ (Table S1). A total of 8 complete putative reductive dehalogenase (RDase) genes were annotated from the draft genome of strain HCBD, exhibiting 23.51–99.60% amino acid sequence similarity with previously characterized RDases (Table S4). Phylogenetic analysis of the 8 rdhA sequences—compared with 40 characterized rdhA genes from OHRB—indicated that rdh (WP_034376989.1), shares 99.60% amino acid identity with KB1-5 (WP_078987038.1), which is an unknown reductive dehalogenase that is expressed in the presence of TCE, cDCE, VC, and 1,2-DCA. Its sequence is closest to rdhA6 in *Dehalococcoides* strain FL2 and has the highest homology (22). It is speculated that it may participate in the initial dehalogenation steps of various chloroethylene compounds. While strain HCBD lacks a complete cobalamin biosynthetic pathway, this phenomenon is common in *Dehalococcoides* (23). A putative Mo-dependent formate dehydrogenase (WP_034376812.1) was identified in the *Dehalococcoides* sp. HCBD draft genome, sharing 100% amino acid sequence identity with formate dehydrogenase of *Dehalococcoides mccartyi* WBC-2 (AOV_98834.1) (24).

### Proteomic detection of putative reductive dehalogenases

A focused analysis of the proteomic data from the HCBD enrichment culture identified eight putative reductive dehalogenases, represented by 29 distinct peptide sequences and 39 peptide-spectrum matches. The identified proteins had sequence coverages ranging from 1.4% to 17.4%, with one to seven unique peptides reported per protein. Notably, WP_034376989.1, the candidate RDase identified by genomic analysis as closely related to KB1-5, was detected with four unique peptides covering 17.4% of its predicted amino acid sequence. WP_437236202.1 was represented by seven unique peptides with 17.2% sequence coverage, whereas WP_015407263.1 was represented by four unique peptides with 9.7% sequence coverage. These results provide protein level evidence for the expression of multiple putative RDases in the enrichment culture, including WP_034376989.1. However, the detection of these proteins alone does not establish their individual contributions to HCBD dechlorination or assign specific enzymes to individual steps in the transformation pathway.

**Figure 5.**
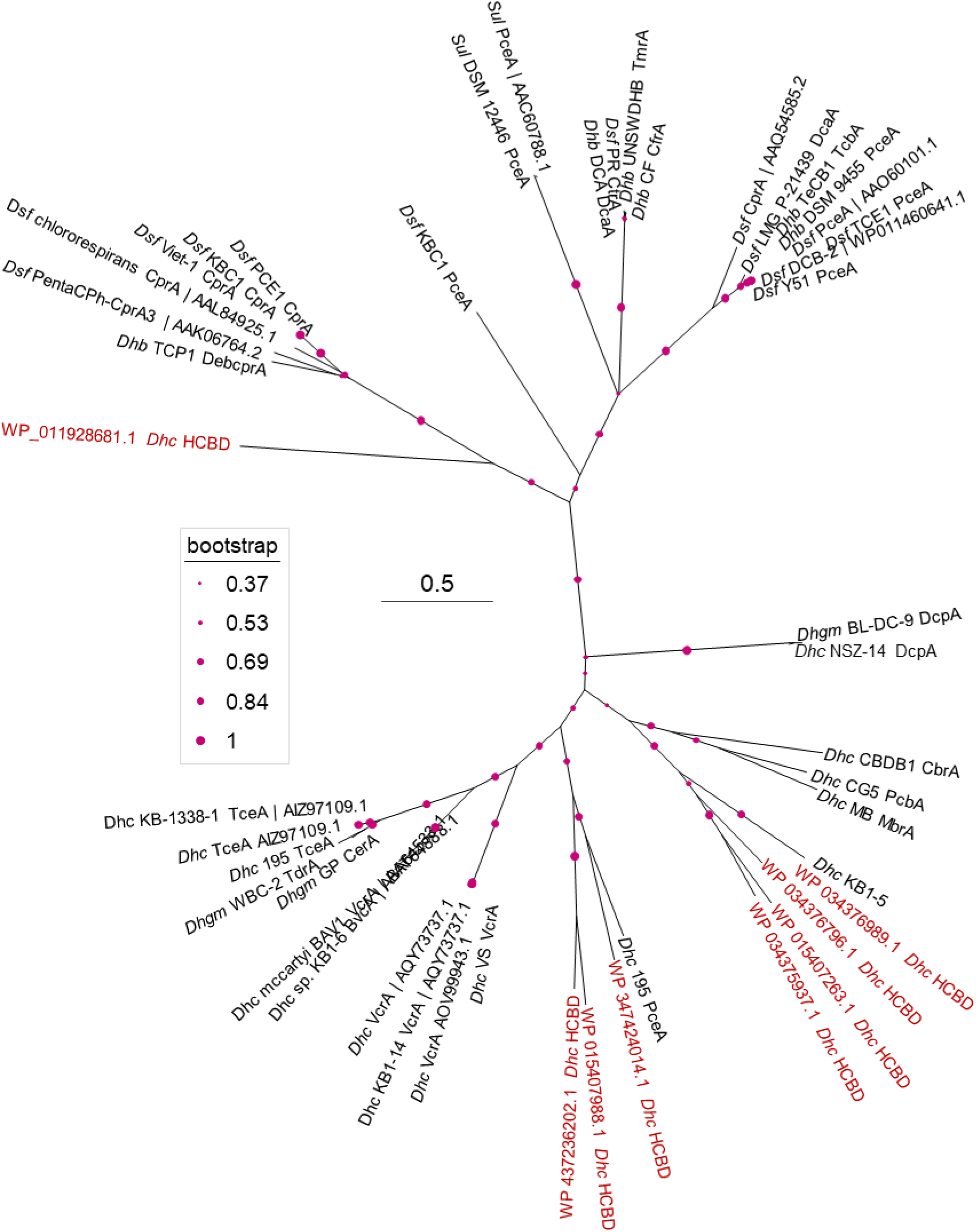
Phylogenetic relationships of reductive dehalogenases (RDases) from strain HCBD and reference OHRB. This neighbor-joining phylogenetic tree compares RDase protein sequences annotated from the *Dhc* sp. HCBD draft genome (red, n = 8). Bootstrap support values are represented by magenta circles, with larger circles indicating higher support.

## Discussion

### HCBD Dechlorination via *Dehalococcoides*

This study demonstrates anaerobic HCBD transformation to less-chlorinated products, including CBD, in *Dehalococcoides*-containing enrichment cultures. The enrichment culture also exhibited complete reductive dechlorination of TCE to ethene, with both transformation processes strongly associated with the activity of *Dehalococcoides*, a genus well known for its exceptional anaerobic dechlorination capacity (21). *Dehalococcoides* species have previously been shown to biotransform a wide range of structurally complex and environmentally persistent organic pollutants, including hexachlorocyclohexane (HCH) isomers, polychlorinated biphenyls (PCBs), and polybrominated diphenyl ethers (PBDEs) (25–27). These compounds are representative of recalcitrant contaminants that pose substantial challenges to conventional remediation strategies due to their chemical stability and resistance to degradation. Although HCBD dechlorination has been reported in enrichment cultures and bioreactor systems (16, 17, 28), the specific microbial taxa responsible for HCBD reductive dechlorination have not been conclusively identified. Previous work demonstrated the co-metabolic transformation of HCBD by *Dehalococcoides* (18), providing early insights into the potential involvement of organohalide-respiring bacteria (OHRB). In contrast to these earlier observations, our results demonstrate that *Dehalococcoides* can utilize HCBD as a terminal electron acceptor in a growth-linked, dechlorination process. This distinction is mechanistically and energetically significant. Co-metabolic transformation relies on fortuitous enzymatic side reactions that do not yield metabolically useful energy, resulting in low and transient transformation rates that depend entirely on alternative growth-supporting substrates (29, 30). In contrast, respiratory reductive dechlorination is directly coupled to energy conservation via electron transport phosphorylation, allowing *Dehalococcoides* to generate a proton motive force and ATP during corrinoid-dependent RDase-mediated dechlorination (31). This energetic gain supports sustained dechlorination activity, active biomass growth, and the induction of substrate-specific RDase genes involved in chlorinated alkene reduction (32). Consequently, once HCBD functions as a bona fide terminal electron acceptor, *Dehalococcoides* can more efficiently channel electron flux toward dechlorination, maintain RDase abundance, and outcompete co-metabolic processes that are inherently inefficient and non-growth-supporting. Overall, the transition from co-metabolism to respiration not only enhances HCBD turnover but also confers a substantial ecological and competitive advantage to *Dehalococcoides* in chlorinated alkene contaminated environments.

### Complex Halogenated Compounds and Microbial Interactions

*Dehalococcoides* may possess a hierarchical electron-acceptor preference, allowing electrons derived from lactate oxidation to be preferentially allocated to energetically favorable and electron-intensive reductive dechlorination pathways, when multiple chlorinated substrates are present. Evidence for such substrate prioritization has been reported previously. For example, *Dehalococcoides mccartyi* exhibits a clear preference for perchloroethene (PCE) and TCE, initiating their dechlorination prior to PCBs, which are transformed only after PCE depletion (33). This behavior suggests that, in the presence of TCE, PCE, and other co-occurring chlorinated compounds like PCBs, *Dehalococcoides* preferentially directs electron flux toward structurally simpler and more energetically accessible substrates like TCE and PCE. Consequently, the transformation of more complex and recalcitrant compounds such as HCBD may be delayed until preferred electron acceptors are depleted. Such substrate-dependent prioritization implies that microbial interactions governing halogenated compound turnover are more complex than previously assumed. For instance, *Dehalococcoides* and *Geobacter* have been shown to exhibit three distinct interaction modes: direct competition, conditional competition, and syntrophic cooperation during the biodegradation of PCE and PCBs (34). These observations further suggest that efficient HCBD dechlorination may likewise depend on coordinated metabolic interactions among multiple microbial taxa. An additional constraint arises from the widespread corrinoid auxotrophy of *Dehalococcoides*. Because *Dehalococcoides* species lack the capacity for de novo corrinoid biosynthesis, these essential cofactors for reductive dehalogenases must be acquired from the surrounding microbial community (35). Strain HCBD exhibits the same corrinoid dependency, underscoring the necessity for syntrophic partnerships and close associations with corrinoid-producing microorganisms in mixed cultures (36).

### Functional Implications of RDases

The putative reductive dehalogenase encoded by the rdhA gene in *Dehalococcoides* sp. strain HCBD shares 99.60% amino acid sequence similarity with the well-characterized dehalogenase from the KB-1 consortium, a mixed culture widely recognized for its effectiveness in TCE bioremediation (37, 38). This exceptionally high degree of sequence similarity is notable, given that strain HCBD was enriched from Xi River sediment in Shenyang, China, a location geographically distinct from the original source of the KB-1 consortium. Amino acid alignment revealed differences at only two positions (I131V and M249L). Because isoleucine, valine, methionine, and leucine are all nonpolar, hydrophobic residues, these substitutions are conservative in nature and are unlikely to disrupt the enzyme’s tertiary structure or catalytic function. Such strong sequence conservation suggests that KB1-5, traditionally associated with respiratory TCE reduction, may possess broader substrate versatility, including the capacity to mediate HCBD dechlorination. This observation has important practical implications. The KB-1 consortium was the first commercially developed microbial culture for in situ bioremediation of chlorinated ethene–contaminated sites (39, 40). The presence of a highly similar reductive dehalogenase in strain HCBD indicates that KB-1 may harbor latent potential to dechlorinate HCBD. By extension, other *Dehalococcoides* strains or consortia carrying KB1-5 homologs may also be capable of transforming HCBD to less chlorinated daughter products. These findings expand the recognized substrate range of KB1-5 enzymes beyond chlorinated ethenes and suggest that existing bioremediation consortia could be evaluated for HCBD degradation with minimal modification. Nevertheless, whether KB1-5 or its close homologs universally confer robust HCBD dechlorination activity remains to be rigorously tested. Future work should include purified enzyme assays, substrate specificity profiling, and controlled bioaugmentation experiments using KB-1 and related cultures to evaluate HCBD transformation efficiency and the extent of complete dechlorination under field-relevant conditions. Such studies will be essential for translating these findings to the remediation of complex sites contaminated with both chlorinated ethenes and HCBD.

### Environmental Implications

The formation of less-chlorinated daughter products during HCBD reductive dechlorination carries important environmental implications that extend beyond demonstrating transformation of the parent persistent organic pollutant. Although sequential dechlorination reduces the degree of halogenation and may increase susceptibility to subsequent biodegradation, the ecotoxicity, persistence, and mobility of partially dechlorinated chlorobutadienes remain poorly constrained and are unlikely to scale linearly with chlorine loss. Consequently, risk reduction cannot be inferred from dechlorination alone, and the environmental fate and toxicity of observed intermediates require targeted evaluation under environmentally relevant conditions.

In this study, the accumulation of CBD highlights a potentially critical bottleneck in HCBD detoxification. These observations suggest that effective remediation of HCBD-contaminated systems may require a treatment train rather than anaerobic bioaugmentation alone. One promising strategy is to couple anaerobic *Dehalococcoides*-mediated dechlorination to lower-chlorinated intermediates with an aerobic polishing step that exploits oxygen-dependent catabolic pathways capable of further transforming or mineralizing chlorinated butadienes. Sequential or spatially segregated redox management approaches such as engineered redox zonation, intermittent aeration, or downstream aerobic bioreactors—may therefore improve remediation endpoints. However, the efficacy of such strategies must be evaluated using quantitative mass balances and toxicity-guided metrics to ensure that daughter-product formation translates into meaningful reductions in environmental hazard. Despite substantial progress over the past two decades in elucidating anaerobic HCBD transformation pathways and the microorganisms involved, HCBD biotransformation remains a critical yet incompletely resolved challenge due to the compound’s environmental persistence, high toxicity, and stringent global regulatory status. This study demonstrates that HCBD can function as a highly favorable electron acceptor under anaerobic conditions, while simultaneously highlighting persistent uncertainties related to metabolic mechanisms, electron-flow constraints, and process robustness under environmentally relevant conditions. Addressing these gaps will require integrative research that combines thermodynamic analyses, genomic and enzymatic characterization, and community-level physiological investigations. Such efforts will be essential for developing predictive frameworks and designing robust, field-applicable bioremediation strategies for HCBD and co-occurring chlorinated contaminants.

## Materials and methods

### Chemicals

Hexachlorobutadiene (CAS number: 87-68-3, HCBD, ≥99 %) was purchased from Aladdin (Shanghai, China). Trichloroethene (CAS number: 79-01-6, TCE, ≥99 %) and cis-1,2-Dichloroethene (CAS number: 156-59-2, cis-DCE, ≥99 %) were ordered from Macklin Co., Ltd. (Shanghai, China). Vinyl chloride (CAS number: 75-01-4, VC, ≥99.5 %) and Ethene (CAS number: 74-85-1, ETH, ≥99.5 %) were obtained from Dalian Special Gases Co., Ltd (Dalian, Liaoning, China). Vitamin B_12_ (≥98 %) was procured from Sigma-Aldrich Chemicals (St. Louis, MO, USA). All other chemicals obtained from Macklin Co., Ltd. (Shanghai, China) or Sigma–Aldrich were of analytical or higher grade.

### Microcosms and transfer cultures

Sampling sites were selected at the Xi River sediments (N 41°37′4″, E 123°3′13″) located in Shenyang City, Liaoning province, China (41). For all cultivation experiments, 120 mL glass serum bottles were utilized, each containing 80 mL of a bicarbonate-buffered, reduced mineral salts medium, prepared according to established protocols (see Text S1 for details) (42). Experimental microcosms were established in triplicate using approximately 5 g of homogenized river sediment slurry, prepared within an anaerobic chamber (Coy Laboratory, Ann Arbor, MI) (43). The vessels were sealed with blue butyl rubber stoppers and secured with aluminum crimps. In the first experimental group, each microcosm was supplemented with 5 mM lactate as both electron donor and carbon source, and Wolin vitamin solution containing 50 μg L⁻¹ vitamin B₁₂. The vitamin solution was filter-sterilized using 0.22 μm polyethersulfone membrane filters (JINTENG, Tianjin, China). HCBD (10 μL) was introduced using dedicated glass microsyringes (Hamilton, Reno, NV, USA). In the second experimental group, each microcosm received 5 mM lactate as both electron donor and carbon source, HCBD (10 μL) and TCE (8 μL) as electron acceptors, and 50 μg L^−1^ Wolin vitamin solution. The initial sediment microcosms amended with HCBD alone or HCBD plus TCE were designated T0-HCBD and T0-HCBD+TCE, respectively. T1, T2, and T3 denote the first, second, and third transfer generations, whereas the suffixes HCBD, HCBD+TCE, and TCE indicate the chlorinated substrates added at the corresponding generation. The HCBD-fed enrichment series was initiated from T0-HCBD and maintained through successive transfers to T1-HCBD, T2-HCBD, and T3-HCBD. T1-HCBD+TCE and T3-HCBD+TCE were established using inocula from T0-HCBD+TCE and T2-HCBD+TCE, respectively. The T3-TCE treatment was inoculated from T2-HCBD+TCE and amended with TCE as the sole added chlorinated substrate. Fresh transfer cultures received 5% v/v inoculum, 5 mM lactate, with 10 μL HCBD and 8 μL TCE, respectively, according to the treatment designation. Control experiments to assess abiotic transformation were conducted using identical preparations subjected to autoclaving prior to incubation. All experimental vessels were maintained under static conditions at 30°C in darkness.

### DNA Extraction and 16S rRNA Gene Amplicon Sequencing

Genomic DNA was extracted from enrichment cultures (1 mL) and 0.25 g sediment using a TIANamp Soil DNA Kit (TIANGEN Biotech, Beijing, China). Prior to extraction, samples underwent vacuum filtration through 0.22-µm mixed cellulose membrane filters (25-mm diameter, JINTENG) (44). DNA quantification was performed with a Qubit 3.0 fluorometer (Invitrogen, Carlsbad, CA, USA). Bacterial community analysis targeted the V3-V4 hypervariable region of the 16S rRNA gene (Table S3). PCR reactions (25 µL) contained template DNA (20 ng), forward and reverse primers (200 nM each), TransStart Taq DNA polymerase (0.5 µL), buffer (2.5 µL), dNTPs (200 µM each), and nuclease-free water. The thermal cycling protocol included initial denaturation (94 °C, 3 min), 24 cycles of denaturation (94 °C, 5 s), annealing (57 °C, 90 s), and extension (72 °C, 10 s), concluding with a final extension (72 °C, 5 min). The MetaVX Library Preparation Kit was employed for library construction, with indexed adapters incorporated through limited cycle PCR. Using 20-30 ng of amplified DNA, sequencing libraries were generated and subsequently analyzed on an Illumina MiSeq platform utilizing 2×250 bp paired-end chemistry at Genewiz Inc (45). Data processing employed Mothur (www.mothur.org) with default parameters. Sequence analysis followed standard MiSeq protocols through the mothur software package (46). The SILVAngs pipeline processed quality-filtered sequences using default settings, with operational taxonomic units (OTUs) clustered at 97 % sequence identity (47).

### Quantitative PCR (qPCR)

*Dehalococcoides* 16S rRNA gene abundance was monitored using primers Dhc16S-q1200F, Dhc16S-q1271R, and probe Dhc16S q1240P (Table S3). The 25 µL reaction mixture contained 12.5 µL of 2x Premix Ex TaqTM master mix (Takara Bio Inc.), primers and probe (0.2 μM each), 50x ROX reference Dye II (0.5 µL), template DNA (2 µL), and nuclease-free water (8.5 µL) (48). A QuantStudioTM 3 Real-Time PCR system (Applied Biosystems, Waltham, MA, USA) was programmed for initial incubation at 50 °C (2 min) and denaturation at 95 °C (10 min), followed by 40 cycles of denaturation (95 °C, 15 s) and annealing/extension (60 °C, 1 min). Quantification relied on a standard curve constructed from serial dilutions of plasmid DNA containing a *Dehalococcoides mccartyi* 195^T^ 16S rRNA gene fragment (GenBank: GCA_000011905.1). The assay demonstrated excellent performance with 102.3% amplification efficiency (slope = −3.268) and strong linearity (R² = 0.999) across a dynamic range of 10^2^-10^8^ gene copies per reaction.

### Metagenomic Analysis

Bacterial biomass was concentrated by centrifugation (13,000 × g, 30 min, 4 °C) prior to genomic DNA extraction using the sodium dodecyl sulfate (SDS) method (49). DNA quality and quantity were assessed via agarose gel electrophoresis and Qubit® 2.0 Fluorometer (Thermo Scientific), respectively. Library preparation and sequencing were performed by Novogene Bioinformatics Technology Co., Ltd. (Beijing, China) on an Illumina NovaSeq 6000 platform. Raw sequence processing employed Readfq (V8, https://github.com/cjfields/readfq) for filtering, with subsequent assembly via the JGI Metagenome Assembly Pipeline (50). Taxonomic characterization occurred at dual levels: metagenomic reads were analyzed using Kaiju v1.7.3 (51), while contigs were binned using MaxBin2 v2.2.4 (52). Quality assessment of metagenome-assembled genomes (MAGs) was conducted with CheckM (53). Taxonomic assignments integrated analyses from GTDB-Tk and the Type Genome Server (TYGS, https://tygs.dsmz.de) (54). Functional annotation of strain HCBD draft genome and other high-quality MAGs was completed using RAST (Rapid Annotations using Subsystems Technology) (55).

### Comparative genomic analysis

Strain HCBD genome was compared with available *Dehalococcoides mccartyi* genomes in GenBank, and digital DNA-DNA hybridization (dDDH) values were calculated via the web-based Genome-to-Genome Distance Calculator 3.0 (GGDC, https://ggdc.dsmz.de/ggdc.php) (56).

### Phylogenetic analyses

MEGA 11.0.9 software was employed for phylogenetic reconstruction. For 16S rRNA gene analysis, sequences were aligned via ClustalW to identify highly homologous GenBank sequences. The neighbor-joining tree incorporated the Tamura-Nei evolutionary model (57) with branch reliability assessed through 1,000 bootstrap replicates (58, 59). Tree visualization utilized the Interactive Tree of Life (iTOL) web platform (https://itol.embl.de/) (60). Separate analyses of reductive dehalogenase (RDase) proteins employed neighbor-joining methodology with the Jones-Taylor-Thornton substitution model (61).

### Analytical methods

Qualitative measurement of HCBD and its transformation products was performed using an ISQ LT purge and trap gas chromatography-mass spectrometry (GC-MS) system. The analysis employed a TRACE 1300 series gas chromatograph coupled to a Single Quadrupole EI-mass spectrometer (Thermo Fisher Scientific) integrated with an OI Analytical 4100 Water/Soil Sample Processor (Xylem Inc., Irvine, CA, USA) and equipped with an Agilent DB-624 column (30 m × 0.25 mm × 1.4 μm). Volatile samples (10 μL gas) were injected with a split ratio of 1:30. The inlet temperature was maintained at 200°C. The column oven temperature was held at 35°C for 6 minutes, then increased to 75°C at a rate of 5°C/min, further raised to 120°C at 10°C/min, and finally elevated to 200°C at 60°C/min, followed by a 5-minute hold. The mass spectral acquisition range was set to m/z 30-300.

The concentrations of TCE, cis-DCE, VC, ETH and Methane were analyzed using an Agilent system consisting of a 7697A GC headspace autosampler coupled to a 7890B gas chromatograph (Agilent Technologies, Santa Clara, CA, USA). The GC was fitted with a DB-624 capillary column (60 m × 0.32 mm × 1.8 μm) and flame ionization detector (FID) (41). Quantification was achieved by comparing peak areas against standard curves, which were generated using known quantities of the target compounds in vessels matching the experimental gas-to-liquid ratios and total volumes. The identification of volatile compounds was conducted using a TRACE 1300 Gas chromatography−mass spectrometry (GC−MS) system linked to a Single Quadrupole EI-mass spectrometer (Thermo Fisher Scientific), integrated with an OI Analytical 4100 Water/Soil Sample Processor (Xylem Inc., Irvine, CA) and fitted with an Agilent DB-624 column. Sample injection (10 μL) employed a 1:30 split ratio with the inlet maintained at 200 °C. The temperature program began at 30 °C (6 min hold), followed by sequential increases to 75 °C (5 °C min^−1^), 120 °C (10 °C min^−1^), and 200 °C (60 °C min^−1^), concluding with a 3.5 min hold.

### Proteomics analysis

Biomass from 1 L of T1-HCBD cultures was harvested by centrifugation at 13,000 × g for 30 min at 4 °C for protein identification. Proteins were extracted using an SDS lysis buffer (4% SDS, 100 mM Tris–HCl, pH 7.6), followed by boiling for 5 min, sonication, and centrifugation at 20,000 × g for 5 min. The supernatants were collected, and proteins were reduced and alkylated with TCEP and CAA at 100 °C for 5 min. Samples were diluted with 50 mM NH_4_HCO_3_ and digested with trypsin (20 ng/µL) at 37 °C for 16–18 h. Following centrifugation at 12,000 × g for 10 min, peptides in the supernatants were desalted using C18 StageTips, vacuum-dried, and reconstituted in 0.1% formic acid. Peptides were separated on an EASY-nLC 1200 system (Thermo Scientific) using a C18 trap column (100 µm × 20 mm, 5 µm particle size) and a C_18_ analytical column (75 µm × 150 mm, 3 µm particle size) at a flow rate of 300 nL min−1. A 60-min gradient from 2% to 100% solvent B (0.1% formic acid in 80% acetonitrile) was applied. Eluted peptides were analyzed using a Q Exactive HF-X mass spectrometer (Thermo Scientific) operated in positive-ion data-dependent acquisition mode. Full MS scans were acquired at a resolution of 60,000 at m/z 200 over an m/z range of 350–1800, with an automatic gain control (AGC) target of 3 × 10^6^ and a maximum injection time of 50 ms. The 20 most intense precursor ions were selected for higher-energy collisional dissociation (HCD) with a normalized collision energy of 28 and an isolation window of 1.6 m/z. MS/MS spectra were acquired at a resolution of 15,000, with an AGC target of 1 × 10^5^ and a maximum injection time of 50 ms. Mass spectrometry data were searched against a reductive dehalogenase (RDase) protein sequence database using MaxQuant. The false discovery rate (FDR) was controlled at ≤1% at both the peptide-spectrum match and protein levels. Protein identification results, including accession numbers, unique peptide counts, and sequence coverage, are summarized in Table 1.

**Table 1.** Proteomic identification of putative reductive dehalogenases in the HCBD enrichment culture.

| Protein accession | Length (aa) | Peptides (n) | Unique peptides (n) | Sequence coverage (%) |
| --- | --- | --- | --- | --- |
| WP_437236202.1 | 494 | 7 | 7 | 17.2 |
| WP_015407263.1 | 497 | 5 | 4 | 9.7 |
| WP_015407988.1 | 505 | 3 | 3 | 12.7 |
| WP_034375937.1 | 475 | 3 | 3 | 5.3 |
| **WP_034376989.1** | **500** | **4** | **4** | **17.4** |
| WP_034376796.1 | 514 | 5 | 4 | 14.8 |
| WP_011928681.1 | 455 | 2 | 2 | 3.5 |
| WP_347424014.1 | 507 | 1 | 1 | 1.4 |

Notes: All entries were annotated as reductive dehalogenases in the search database. Unique peptides are defined relative to the database used for identification. Protein lengths and sequence coverage refer to the database entries; WP\_437236202.1 is annotated as a partial sequence. Shared peptides may be assigned to more than one protein, so peptide counts are not additive. The dataset comprised 29 distinct peptide sequences and 39 peptide-spectrum matches across eight protein groups. Boldface highlights WP\_034376989.1, the candidate discussed in the genomic analysis. aa, amino acids.

## Data availability

The raw metagenomic sequencing reads of the HCBD-dechlorinating enrichment culture, used for microbial community profiling and reconstruction of the *Dehalococcoides* sp. strain HCBD genome, have been deposited in the NCBI Sequence Read Archive (SRA) under accession number SRR34078029, with the associated BioSample accession number SAMN49504897.

## Acknowledgements

This work was supported by the National Natural Science Foundation of China (Grant No. 42177220), the Intergovernmental International Cooperation in Science and Technology Innovation of National Key Research and Development Project of China (Grant No. 2023YFE0122000), the Shanghai Frontiers Science Center of Polar Science (Grant No. SOO2025-01), the Open Funding of State Key Laboratory of Coal Mine Disaster Prevention and Control (Grant No. 2024SKLKF03), the International Science and Technology Cooperation Project of Department of Science and Technology of Liaoning Province (Grant No. 2024JH2/101900003), and the Partnership Network for Future Innovation Special Program of Chinese Academy of Sciences (Grant No. 064GJHZ2022040FN).

## References

1. Rossberg M, Lendle W, Pfleiderer G, Tögel A, Dreher E-L, Langer E, Rassaerts H, Kleinschmidt P, Strack H, Cook R, Beck U, Lipper K-A, Torkelson TR, Löser E, Beutel KK, Mann T. 2006. Chlorinated Hydrocarbons, Ullmann’s Encyclopedia of Industrial Chemistry 10.1002/14356007.a06_233.pub2.

2. Marshall KA, Pottenger LH. 2016. Chlorocarbons and Chlorohydrocarbons, p 1-29, Kirk-Othmer Encyclopedia of Chemical Technology 10.1002/0471238961.1921182218050504.a01.pub3.

3. Act CEP. 1993. Priority substances list assessment report. Environment Canada and Health Canada, Ottawa, Ontario:1–56.

4. Zhao C, Yang L, Sun Y, Chen C, Huang Z, Yang Q, Yun J, Habib A, Liu G, Zheng M, Jiang G. 2024. Atmospheric emissions of hexachlorobutadiene in fine particulate matter from industrial sources. Nature Communications 15:4737.

5. Tang Z, Huang Q, Cheng J, Qu D, Yang Y, Guo W. 2014. Distribution and accumulation of hexachlorobutadiene in soils and terrestrial organisms from an agricultural area, East China. Ecotoxicology and Environmental Safety 108:329–334.

6. Lu Y, Chen Z-F, Chen Y-J, Xu Y-Z, Chen Y, Dai X, Yao L, Qi Z, Cai Z. 2021. Distribution and risk assessment of hexachlorobutadiene, pentachloroanisole, and chlorobenzenes in sediment and wild fish from a region affected by industrial and agricultural activities in South China. Journal of hazardous materials 417:126002.

7. Yang M, Mao H, Li H, Yang F, Cao F, Yan W. 2023. Quantifying Concentrations and Emissions of Hexachlorobutadiene - A New Atmospheric Persistent Organic Pollutant in northern China. Environmental Research 216:114139.

8. Mackay D, Shiu W-Y, Lee SC. 2006. Handbook of physical-chemical properties and environmental fate for organic chemicals. CRC press.

9. Chen X, Luo Q, Wang D, Gao J, Wei Z, Wang Z, Zhou H, Mazumder A. 2015. Simultaneous assessments of occurrence, ecological, human health, and organoleptic hazards for 77 VOCs in typical drinking water sources from 5 major river basins, China. Environmental pollution 206:64–72.

10. Zhang H, Wang Y, Sun C, Yu M, Gao Y, Wang T, Liu J, Jiang G. 2014. Levels and distributions of hexachlorobutadiene and three chlorobenzenes in biosolids from wastewater treatment plants and in soils within and surrounding a chemical plant in China. Environmental science & technology 48:1525–1531.

11. Kong Q, Wang Y, Yang X. 2020. A Review on Hexachloro-1,3-butadiene (HCBD): Sources, Occurrence, Toxicity and Transformation. Bulletin of Environmental Contamination and Toxicology 104:1–7.

12. Li X, Wang M, Hou M, Su G, Sun B, Hua Y, Pang J, Meng J, Shi B, Li Q. 2025. Current status and strategies for controlling hexachlorobutadiene from multiple perspectives of emission, occurrence, and disposal. Environmental Research 268:120760.

13. UNEP. Guidance on the global monitoring plan for persistent organic pollutants, p. In (ed), Secretariat of the Stockholm Convention on Persistent Organic Pollutants …,

14. Li MT, Hao LL, Sheng LX, Xu JB. 2008. Identification and degradation characterization of hexachlorobutadiene degrading strain Serratia marcescens HL1. Bioresource Technology 99:6878–6884.

15. Cord-Ruwisch R, James D, Charles W. 2009. The use of redox potential to monitor biochemical HCBD dechlorination. Journal of biotechnology 142:151–156.

16. Bosma TNP, Cottaar FHM, Posthumus MA, Teunis CJ, van Veldhuizen A, Schraa G, Zehnder AJB. 1994. Comparison of Reductive Dechlorination of Hexachloro-1,3-butadiene in Rhine Sediment and Model Systems with Hydroxocobalamin. Environmental Science & Technology 28:1124–1128.

17. Booker RS, Pavlostathis SG. 2000. Microbial reductive dechlorination of hexachloro-1,3-butadiene in a methanogenic enrichment culture. Water Research 34:4437–4445.

18. Shen R, Zhang S, Liang Z, Mai B, Wang S. 2022. Mechanistic insight into co-metabolic dechlorination of hexachloro-1,3-butadiene in Dehalococcoides. Water Research 220:118725.

19. James DL. 2009. Biochemical dechlorination of Hexachloro-1, 3-butadiene. Murdoch University.

20. Luo Y, Liu G. 2024. Chemical mechanisms of hexachlorobutadiene reactions in the environment. Environmental Pollution 361:124893.

21. Löffler FE, Yan J, Ritalahti KM, Adrian L, Edwards EA, Konstantinidis KT, Müller JA, Fullerton H, Zinder SH, Spormann AM. 2013. Dehalococcoides mccartyi gen. nov., sp. nov., obligately organohalide-respiring anaerobic bacteria relevant to halogen cycling and bioremediation, belong to a novel bacterial class, Dehalococcoidia classis nov., order Dehalococcoidales ord. nov. and family Dehalococcoidaceae fam. nov., within the phylum Chloroflexi. International Journal of Systematic and Evolutionary Microbiology 63:625–635.

22. Waller Alison S, Krajmalnik-Brown R, Löffler Frank E, Edwards Elizabeth A. 2005. Multiple Reductive-Dehalogenase-Homologous Genes Are Simultaneously Transcribed during Dechlorination by Dehalococcoides-Containing Cultures. Applied and Environmental Microbiology 71:8257–8264.

23. Seshadri R, Adrian L, Fouts DE, Eisen JA, Phillippy AM, Methe BA, Ward NL, Nelson WC, Deboy RT, Khouri HM, Kolonay JF, Dodson RJ, Daugherty SC, Brinkac LM, Sullivan SA, Madupu R, Nelson KE, Kang KH, Impraim M, Tran K, Robinson JM, Forberger HA, Fraser CM, Zinder SH, Heidelberg JF. 2005. Genome Sequence of the PCE-Dechlorinating Bacterium Dehalococcoides ethenogenes. Science 307:105–108.

24. Molenda O, Tang S, Edwards EA. 2016. Complete Genome Sequence of Dehalococcoides mccartyi Strain WBC-2, Capable of Anaerobic Reductive Dechlorination of Vinyl Chloride. Genome Announcements 4:10.1128/genomea.01375-16.

25. Bashir S, Kuntze K, Vogt C, Nijenhuis I. 2018. Anaerobic biotransformation of hexachlorocyclohexane isomers by Dehalococcoides species and an enrichment culture. Biodegradation 29:409–418.

26. Chen C, Xu G, Rogers MJ, He J. 2024. Metabolic Synergy of Dehalococcoides Populations Leading to Greater Reductive Dechlorination of Polychlorinated Biphenyls. Environmental Science & Technology 58:2384–2392.

27. Zhao S, Ding C, Xu G, Rogers MJ, Ramaswamy R, He J. 2022. Diversity of organohalide respiring bacteria and reductive dehalogenases that detoxify polybrominated diphenyl ethers in E-waste recycling sites. The ISME Journal 16:2123–2131.

28. Boucquey JB, Renard P, Amerlynck P, Filho PM, Agathos S, Naveau H, Nyns EJ. 1995. High-rate continuous biodegradation of concentrated chlorinated aliphatics by a durable enrichment of methanogenic origin under carrier-dependent conditions. Biotechnology and bioengineering 47:298–307.

29. Smidt H, de Vos WM. 2004. Anaerobic Microbial Dehalogenation. Annual Review of Microbiology 58:43–73.

30. Chang H-L, Alvarez-Cohen L. 1995. Model for the Cometabolic Biodegradation of Chlorinated Organics. Environmental Science & Technology 29:2357–2367.

31. Müller JA, Rosner BM, Abendroth Gv, Meshulam-Simon G, McCarty PL, Spormann AM. 2004. Molecular Identification of the Catabolic Vinyl Chloride Reductase from Dehalococcoides sp. Strain VS and Its Environmental Distribution. Applied and Environmental Microbiology 70:4880–4888.

32. Krajmalnik-Brown R, Hölscher T, Thomson IN, Saunders FM, Ritalahti KM, Löffler FE. 2004. Genetic Identification of a Putative Vinyl Chloride Reductase in Dehalococcoides sp. Strain BAV1. Applied and Environmental Microbiology 70:6347–6351.

33. Chen C, He J. 2018. Strategy for the Rapid Dechlorination of Polychlorinated Biphenyls (PCBs) by Dehalococcoides mccartyi Strains. Environmental Science & Technology 52:13854–13862.

34. Liang Y, Lu Q, Liang Z, Liu X, Fang W, Liang D, Kuang J, Qiu R, He Z, Wang S. 2021. Substrate-dependent competition and cooperation relationships between Geobacter and Dehalococcoides for their organohalide respiration. ISME Communications 1:23.

35. Türkowsky D, Jehmlich N, Diekert G, Adrian L, von Bergen M, Goris T. 2018. An integrative overview of genomic, transcriptomic and proteomic analyses in organohalide respiration research. FEMS Microbiology Ecology 94.

36. Kruse S, Türkowsky D, Birkigt J, Matturro B, Franke S, Jehmlich N, von Bergen M, Westermann M, Rossetti S, Nijenhuis I, Adrian L, Diekert G, Goris T. 2021. Interspecies metabolite transfer and aggregate formation in a co-culture of Dehalococcoides and Sulfurospirillum dehalogenating tetrachloroethene to ethene. The ISME Journal 15:1794–1809.

37. Saiyari DM, Chuang H-P, Senoro DB, Lin T-F, Whang L-M, Chiu Y-T, Chen Y-H. 2018. A review in the current developments of genus Dehalococcoides, its consortia and kinetics for bioremediation options of contaminated groundwater. Sustainable Environment Research 28:149–157.

38. Krajmalnik-Brown R, Sung Y, Ritalahti KM, Michael Saunders F, Löffler FE. 2007. Environmental distribution of the trichloroethene reductive dehalogenase gene (tceA) suggests lateral gene transfer among Dehalococcoides. FEMS Microbiology Ecology 59:206–214.

39. Major DW, McMaster ML, Cox EE, Edwards EA, Dworatzek SM, Hendrickson ER, Starr MG, Payne JA, Buonamici LW. 2002. Field demonstration of successful bioaugmentation to achieve dechlorination of tetrachloroethene to ethene. Environmental Science & Technology 36:5106–5116.

40. Lyon DY, Vogel TM. 2013. Bioaugmentation for Groundwater Remediation: an Overview, p 1–37. In Stroo HF, Leeson A, Ward CH (ed), Bioaugmentation for Groundwater Remediation doi:10.1007/978-1-4614-4115-1_1. Springer New York, New York, NY.

41. Li X, Wang J, Yang Y, Jin H, Wang H, Zhang Z, Li X, Cui Y, Wang X, Yan J. 2024. Sequential Reductive Dechlorination of Triclosan by Sediment Microbiota Harboring Organohalide-Respiring Dehalococcoidia. Environmental Science & Technology 58:14843–14854.

42. Löffler FE, Sanford RA, Ritalahti KM. 2005. Enrichment, cultivation, and detection of reductively dechlorinating bacteria. Methods Enzymol 397:77–111.

43. Wang H, Jin H, Wang J, Wang X, Li X, Yan J, Yang Y. 2024. Dehalogenimonas Strain W from Estuarine Sediments Dechlorinates 1,2-Dichloroethane under Elevated Salinity. Environmental Science & Technology doi:10.1021/acs.est.4c08999.

44. Yang Y, Cápiro NL, Marcet TF, Yan J, Pennell KD, Löffler FE. 2017. Organohalide Respiration with Chlorinated Ethenes under Low pH Conditions. Environmental Science & Technology 51:8579–8588.

45. Kozich JJ, Westcott SL, Baxter NT, Highlander SK, Schloss PD. 2013. Development of a Dual-Index Sequencing Strategy and Curation Pipeline for Analyzing Amplicon Sequence Data on the MiSeq Illumina Sequencing Platform. Applied and Environmental Microbiology 79:5112–5120.

46. Hiltemann SD, Boers SA, van der Spek PJ, Jansen R, Hays JP, Stubbs AP. 2019. Galaxy mothur Toolset (GmT): a user-friendly application for 16S rRNA gene sequencing analysis using mothur. GigaScience 8:giy166.

47. Quast C, Pruesse E, Yilmaz P, Gerken J, Schweer T, Yarza P, Peplies J, Glöckner FO. 2012. The SILVA ribosomal RNA gene database project: improved data processing and web-based tools. Nucleic Acids Research 41:D590–D596.

48. Jiang L, Yang Y, Jin H, Wang H, Swift CM, Xie Y, Schubert T, Löffler FE, Yan J. 2022. Geobacter sp. Strain IAE Dihaloeliminates 1,1,2-Trichloroethane and 1,2-Dichloroethane. Environmental Science & Technology.

49. Wang H, Wang X, Huang S, Yang S, Liao H, Wang X, Jin H, Wang J, Li X, Yan J, Yang Y. Complete genome sequence of “Candidatus Dehalogenimonas loeffleri”strain W, a highly salt-tolerant chlorinated alkane-dechlorinating bacterium isolated from estuarine sediments. Microbiology Resource Announcements 0:e00319–24.

50. Arkin AP, Cottingham RW, Henry CS, Harris NL, Stevens RL, Maslov S, Dehal P, Ware D, Perez F, Canon S. 2018. KBase: the United States department of energy systems biology knowledgebase. Nature Biotechnology 36:566–569.

51. Menzel P, Ng KL, Krogh A. 2016. Fast and sensitive taxonomic classification for metagenomics with Kaiju. Nature Communications 7:11257.

52. Wu Y-W, Simmons BA, Singer SW. 2016. MaxBin 2.0: an automated binning algorithm to recover genomes from multiple metagenomic datasets. Bioinformatics 32:605–607.

53. Parks DH, Imelfort M, Skennerton CT, Hugenholtz P, Tyson GW. 2015. CheckM: assessing the quality of microbial genomes recovered from isolates, single cells, and metagenomes. Genome Research 25:1043–1055.

54. Meier-Kolthoff JP, Göker M. 2019. TYGS is an automated high-throughput platform for state-of-the-art genome-based taxonomy. Nature Communications 10:2182.

55. Overbeek RA, Olson R, Pusch GD, Olsen GJ, Davis JJ, Disz T, Edwards RA, Gerdes SY, Parrello BD, Shukla M, Vonstein V, Wattam AR, Xia F, Stevens RL. 2013. The SEED and the Rapid Annotation of microbial genomes using Subsystems Technology (RAST). Nucleic Acids Research 42:D206–D214.

56. Meier-Kolthoff JP, Carbasse JS, Peinado-Olarte RL, Göker M. 2022. TYGS and LPSN: a database tandem for fast and reliable genome-based classification and nomenclature of prokaryotes. Nucleic Acids Research 50:D801–D807.

57. Tamura K, Nei M. 1993. Estimation of the number of nucleotide substitutions in the control region of mitochondrial DNA in humans and chimpanzees. Molecular Biology and Evolution 10:512–526.

58. Jin H, Li X, Wang H, Cápiro NL, Li X, Löffler FE, Yan J, Yang Y. 2022. Anaerobic Biohydrogenation of Isoprene by Acetobacterium wieringae Strain Y. mBio 13:e02086–22.

59. Saitou N, Nei M. 1987. The neighbor-joining method: a new method for reconstructing phylogenetic trees. Molecular Biology and Evolution 4:406–425.

60. Letunic I, Bork P. 2006. Interactive Tree Of Life (iTOL): an online tool for phylogenetic tree display and annotation. Bioinformatics 23:127–128.

61. Jones DT, Taylor WR, Thornton JM. 1992. The rapid generation of mutation data matrices from protein sequences. Computer Applications in the Biosciences : CABIOS 8 3:275–82.

